# α-Synuclein oligomers form by secondary nucleation

**DOI:** 10.1101/2023.05.28.542651

**Authors:** Catherine K Xu, Georg Meisl, Ewa Andrzejewska, Georg Krainer, Alexander J Dear, Marta Castellana Cruz, Soma Turi, Raphael Jacquat, William E Arter, Michele Vendruscolo, Sara Linse, Tuomas PJ Knowles

## Abstract

Oligomeric species arising during aggregation of α-synuclein are proposed to be a major source of toxicity in Parkinson’s disease, and thus a major potential drug target. However, their mechanism of formation and role in aggregation are largely unresolved. Here we first show that, at physiological pH, α-synuclein aggregates by secondary nucleation, rather than fragmentation, and that this process is enhanced by agitation. Moreover, using a combination of single molecule and bulk level techniques, we identify secondary nucleation on the surfaces of existing fibrils, rather than formation directly from monomers, as the dominant source of oligomers. Our results highlight secondary nucleation as not only the key source of oligomers, but also the main mechanism of aggregate formation, and show that these processes take place under physiologically relevant conditions.

## Introduction

The process of protein aggregation is associated with over 50 human disorders, including Alzheimer’s and Parkinson’s diseases (PD) [1]. In PD, aggregates of the 14 kDa protein α-synuclein are the major component of Lewy bodies and neurites, which emerge as the pathological hallmarks of the disease [2]. In addition to its abundance in the characteristic amyloid deposits in PD, α-synuclein is further implicated in PD disease development by the finding that duplications and triplications of the WT α-synuclein gene, as well as a number of single-point mutations, are associated with familial cases of PD [3, 4].

While deposits of fibrillar protein are hallmarks of protein aggregation diseases, oligomeric intermediates are proposed to be the major source of toxicity [2, 5, 6, 1, 7]. Moreover, determining oligomer dynamics is crucial for the elucidation of aggregation mechanisms [8, 9]. Oligomers are nevertheless relatively poorly characterised, due to challenges in their analysis that render them invisible to most conventional biophysical techniques, namely their low abundance, low stability, and high degree of heterogeneity [10, 11, 12, 13, 8, 9, 14, 7].

Oligomer dynamics during amyloid aggregation the Alzheimer’s disease-associated Aβ peptide have been used to determine the molecular pathways of fibril formation [15]. Oligomers were isolated by size exclusion chromatography, providing an upper limit on oligomer concentrations since interactions with the solid matrix may perturb the system. In the case of α-synuclein, several studies of oligomer dynamics have employed single molecule FRET (smFRET) experiments, which have observed the interconversion oligomers with different FRET efficiencies [10, 16, 17, 18]. However, such single molecule level measurements require extremely high dilution factors of up to 10^6^ to reach nanomolar or even picomolar measurement concentrations, which can destabilise transient oligomers [19].

Here, we exploit the power of microfluidic free-flow electrophoresis (µFFE) at the single molecule level to study the dynamics of α-synuclein oligomers under native-like conditions [20]. Combined with bulk method and chemical kinetics, this approach allows us to quantitatively determine the molecular mechanism of α-synuclein fibril formation under physiologically relevant conditions (Figure 1).

**Figure 1:**
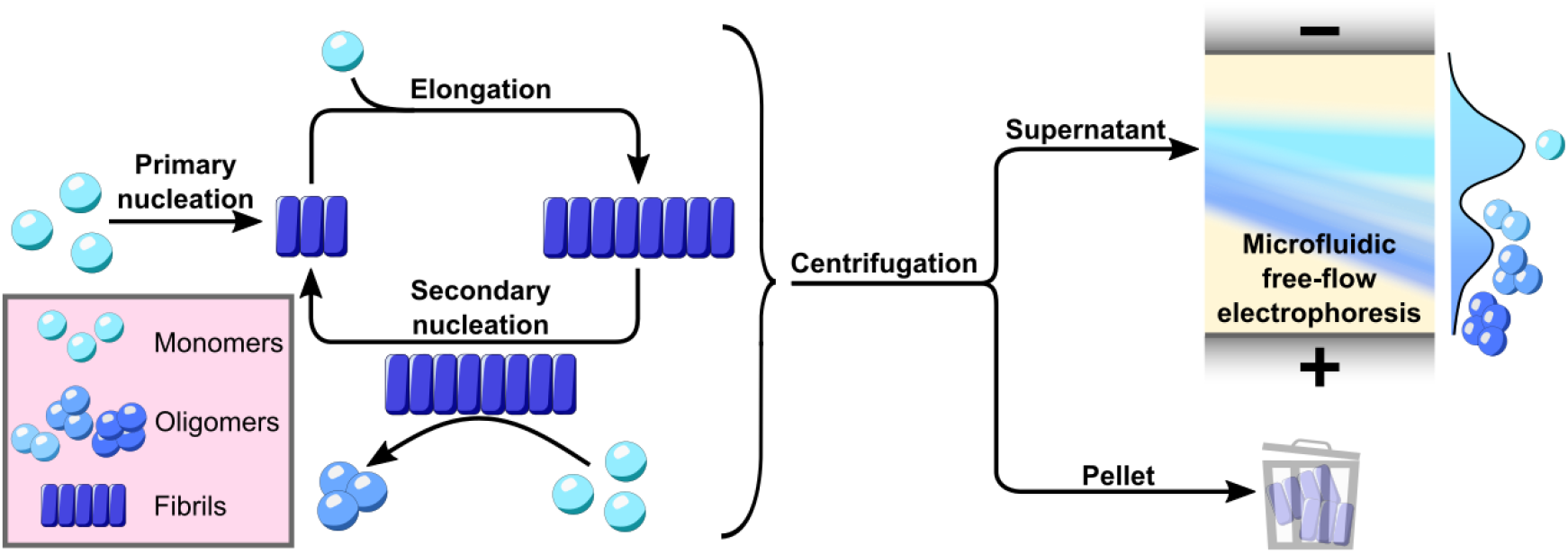
Using microfluidic free-flow electrophoresis at the single molecule level, we were able to fractionate α-synuclein aggregation mixtures and determine oligomer dynamics. We then used chemical kinetics to determine that secondary nucleation on fibril surfaces is the dominant mechanism of both α-synuclein oligomers and new fibrils.

## Results

### *α*-Synuclein aggregation occurs via secondary pathways

Despite its heavy implication in Parkinson’s disease, the aggregation of α-synuclein remains relatively poorly characterised. Detailed investigations into the individual microscopic processes have been carried out under diverse reaction conditions which deviate from physiological pH and/or ionic strength; primary nucleation induced by synthetic lipids which do not represent those abundant in biological membranes, secondary nucleation at moderate ionic strength at mildly acidic pH, and elongation at moderate ionic strength at neutral pH [21, 22, 23]. From such studies, we now qualitatively understand certain aspects of the aggregation mechanism but have so far failed to formulate a quantitative model that can account for the experimental data. In order to unify the reaction steps into one complete mechanistic description of fibril formation from α-synuclein, we established experimental conditions for the reproducible aggregation of α-synuclein at neutral pH (Figures 2 and S7) [24].

**Figure 2:**
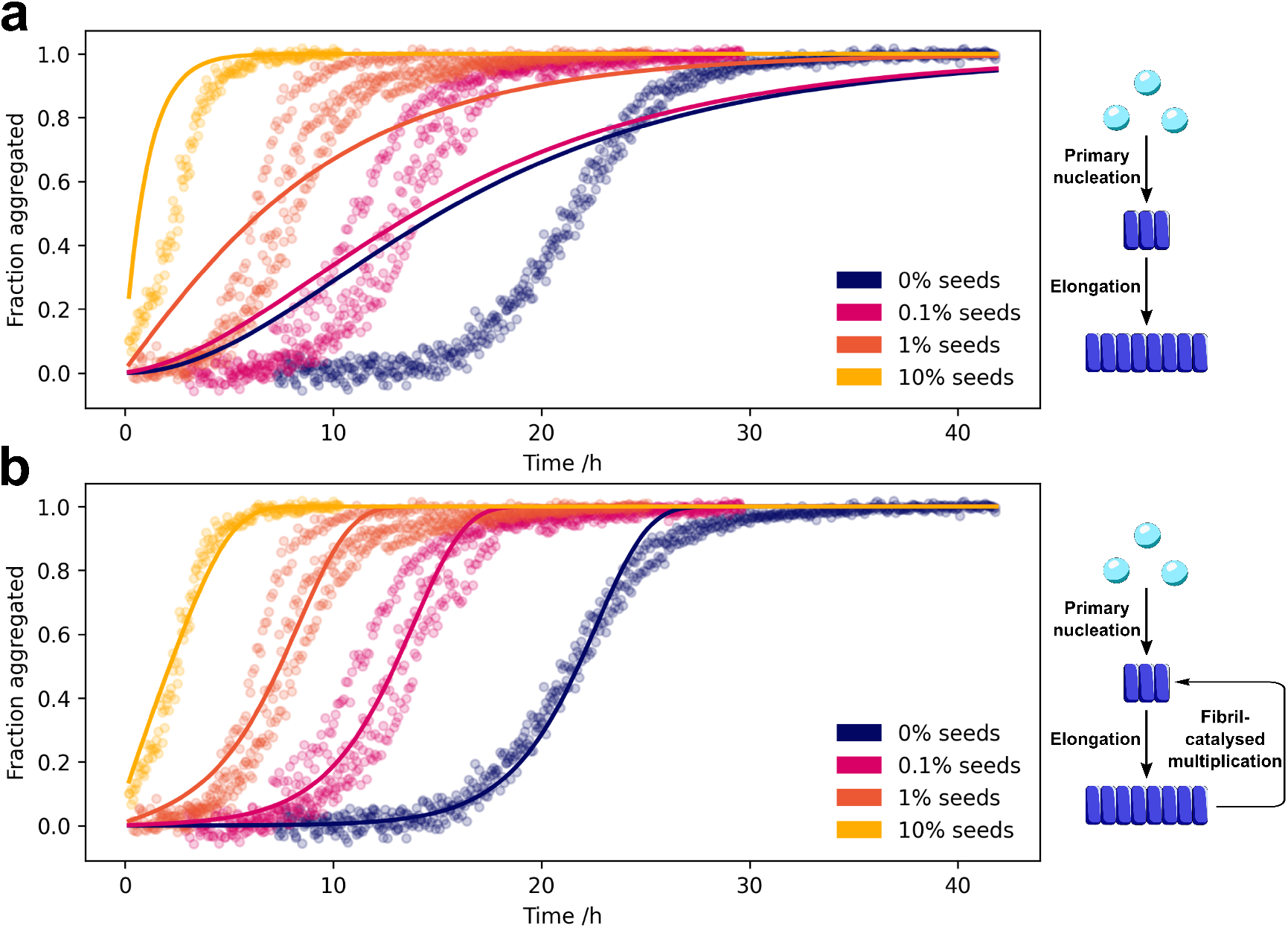
Secondary processes are involved in the aggregation of α-synuclein. The aggregation kinetics of AlexaFluor-488-labelled α-synuclein was followed by aggregation-induced quenching of the AlexaFluor-488 dye (Figure S6). The dependence of α-synuclein aggregation kinetics on the absence and presence of varying concentration of fibrillar seeds. Data were fitted to kinetic models in the absence (**a**) and presence (**b**) of secondary processes. The data are only consistent with a model that includes fibril-catalysed formation of new fibrils.

We investigated the fibril formation kinetics of α-synuclein in the absence and presence of varying concentrations of fibrillar seeds. Upon the addition of fibrillar seeds, the lag phase and aggregation half-time (*t*_1/2_) decreased in a seed concentration-dependent manner. This behaviour is highly characteristic of the presence of secondary processes, whereby existing fibrils catalyse the formation of further fibrils [25, 26]. In contrast, in the absence of secondary processes, the addition of seed fibrils would not significantly alter the aggregation kinetics. Moreover, fitting our data globally to kinetic models, we found that the aggregation kinetics are inconsistent with a mechanism which does not include secondary processes; α-synuclein therefore aggregates via secondary processes [27].

### Secondary nucleation is the dominant mechanism of fibril formation

The dependence of aggregation kinetics on protein concentration can be used to infer the molecular mechanisms of fibril formation [27, 25]. The concentration dependence of the unseeded aggregation rates, with a scaling exponent of −0.5, is consistent with fibril formation mechanisms of both fragmentation and saturated secondary nucleation. In the former case, the number of fibrils increases by the fragmentation of existing fibrils. In the latter case, monomers bind to fibril surfaces, and their subsequent conversion to aggregates and release into solution is the rate limiting step [23, 25]. In order to determine which of the two mechanisms is dominant, we investigated the fibril fragmentation rate. We obtained the fibril length distributions by transmission electron microscopy (TEM) imaging of aggregation mixtures throughout the plateau phase, finding a decrease in the mean fibril length over time (Figures 3, S8, and S9).

**Figure 3:**
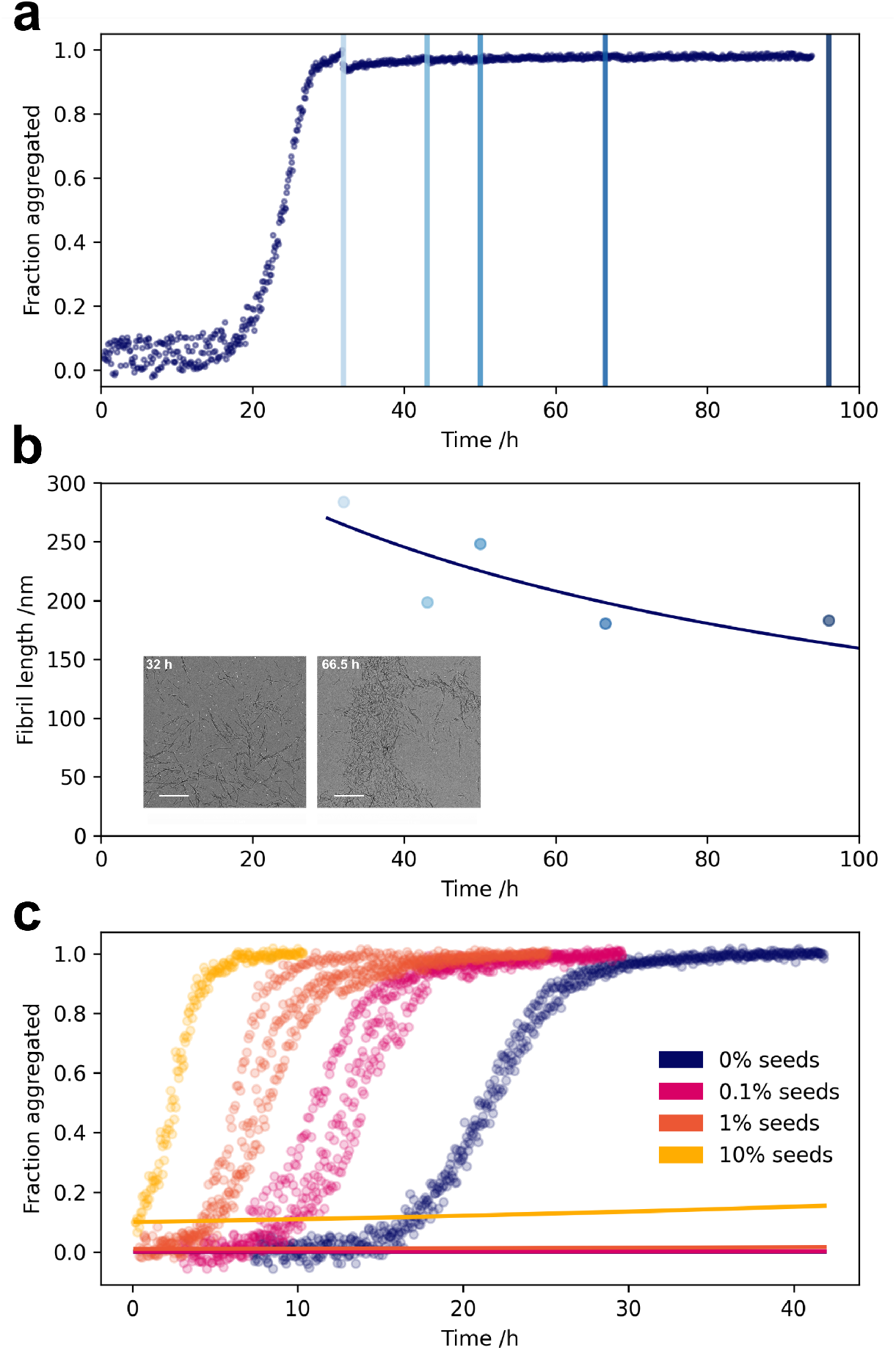
Fragmentation is not the dominant mechanism of α-synuclein fibril formation. Fibrils were withdrawn from aggregation reactions at the indicated timepoints in the plateau phase of the aggregation mixture (**a**) and imaged by TEM (**b**, inset). The mean lengths of fibrils were fitted to kinetic models to determine the fragmentation rate (**b**). The kinetic data were then fitted with fragmentation as the mechanism of fibril amplification, with the fragmentation rate constant fixed at the value determined from the time dependence of the fibril lengths (**c**).

From fitting an analytical expression for the average length (derivation see SI) to the fibril lengths measured by TEM (Figure 3), we obtain an upper bound on the rate of fibril accumulation due to fragmentation *κ*(frag) = 0.01 h^−1^. The aim is to now compare this value, obtained from measurements of fibril lengths, to the actual rate of aggregate formation, *κ*, measured by fluorescence and thus judge if fragmentation alone is sufficient to account for the speed of fibril accumulation seen in the aggregation reaction.

Fitting the aggregation kinetics (Figure 3) yields a very clear result: *κ* = 0.4 h^−1^, i.e. 40-fold faster than would be expected based on fragmentation alone. We therefore conclude that fragmentation is only a minor contributor to the rate of fibril formation, and that secondary nucleation is the dominant mechanism of α-synuclein fibril formation.

### Oligomers form by secondary nucleation on fibril surfaces

In order to elucidate further details of the secondary nucleation process, we investigated α-synuclein oligomer dynamics during aggregation. We previously demonstrated the ability of µFFE to fractionate complex aggregation mixtures and resolve oligomeric subpopulations according to their electrophoretic mobilities, a function of radius and charge (Figure S10) [20]. We have further demonstrated its extension to single molecule spectroscopy to maximise information yield [28]. Here, we employ µFFE at the single molecule level to monitor oligomer mass concentrations during α-synuclein aggregation to yield insights into oligomer dynamics.

A key feature of this approach in studying oligomers is its minimal perturbation of the reaction system. The sample under study is rapidly diluted and fractionated in solution just a few milliseconds prior to measurement, a timescale in which the sample composition is unlikely to change. This contrasts with more traditional single molecule approaches requiring the million-fold dilution of samples, or size exclusion chromatography, where samples interact differentially with a solid matrix on a timescale of minutes to hours [10, 19, 15]. Moreover, the method is agnostic to oligomer structure, as oligomers are detected by their intrinsic fluorescent label, in contrast to antibody-based methods such as ELISA [29, 30, 31, 32]. The reaction mixture is therefore minimally perturbed for the measurement; the attachment of the AlexaFluor-488 fluorophore at position 122 did not affect the aggregation kinetics compared to the WT protein (Figure S5), likely due to its location outside of the fibril core [33].

Using out microfluidic approach, we observed a maximum in oligomer mass concentration slightly before the half-time. Moreover, the peak in oligomer mass concentration in the seeded reaction was similarly located close to the half-time, pointing to secondary nucleation as the dominant mechanism of oligomer formation (Figure 4). Due to the high concentration of monomer and thus background noise, we used a simple photon count thresholding approach to estimate the oligomer concentration. However, the non-uniformity of the confocal spot laser intensity means that the estimated oligomer concentrations may be on the order of ten- or even thousand-fold smaller than the actual concentrations (see SI for details). Using the relative oligomer concentrations, we were able to use chemical kinetics to determine their stability and relative formation rate [8] (see SI for details of the kinetic model). Secondary nucleation is therefore not only responsible for the formation of new fibrils, but is also the main source of oligomers.

**Figure 4:**
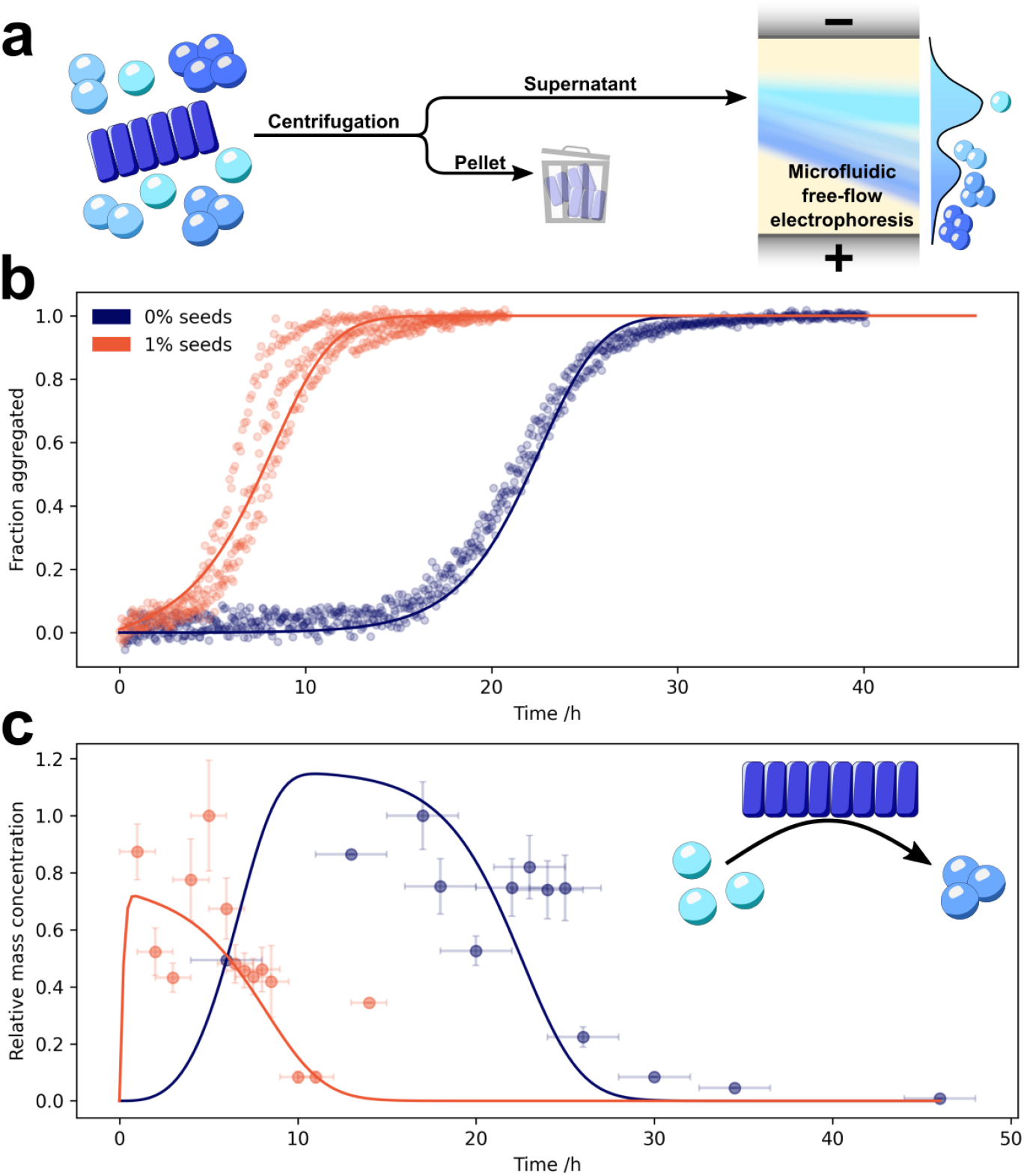
α-Synuclein oligomers form by secondary nucleation. Aggregation mixtures at various timepoints throughout the reaction were centrifuged (21,130 rcf, 10 mins, 20 °C) to remove large fibrillar aggregates, and the oligomer content of the resulting supernatant studied by µFFE at the single molecule level (**a**). Kinetics of fibrillar α-synuclein in unseeded (blue) and seeded (red, 1% seeds) aggregation reactions, measured by fluorescence quenching, are shown alongside the fitted model (**b**). The relative oligomer mass concentrations were determined (**c**) and fitted to a model in which oligomers can form via both primary nucleation from monomers and secondary nucleation on fibril surfaces (model details in SI). X-axis error bars represent the time range over which data were averaged, which corresponds to the standard deviation of the aggregation half times. Where present, y-axis error bars represent the standard error of the mean oligomer mass concentrations; points wihout y-axis error bars represent a single sample.

### Oligomers form under quiescent, native-like conditions

We next investigated the role of shaking in α-synuclein aggregation. In the absence of shaking, aggregation proceeded at a much lower rate, demonstrating that shaking increases the rate of aggregation. However, the addition of seeds drastically reduced the *t*_1/2_, indicating that secondary processes still take place under quiescent conditions. In order to determine which microscopic process/es are catalysed by shaking, we studied the oligomer content of quiescent, seeded aggregation reactions using µFFE at the single molecule level before and after moderate shaking (10 minutes, 200 rpm). Prior to shaking, very few oligomers were observed, but the concentration of oligomers increased by more than a factor of three following moderate shaking (Figure 5). The post-shaking concentration was also higher than the oligomer concentration during the aggregation plateau phase under constant shaking at the same speed (200 rpm), demonstrating that the majority of these oligomers did not arise through fragmentation of fibrils induced by the shaking. Given that the conversion and dissociation steps are rate-limiting in this system, these must be the processes affected by agitation. From a mechanical perspective, the dissociation of oligomers from fibrils is more likely to be catalysed by shaking. We therefore conclude that α-synuclein oligomers form by secondary nucleation under quiescent conditions at neutral pH, and that shaking accelerates their formation, likely by facilitating their dissociation from fibril surfaces.

**Figure 5:**
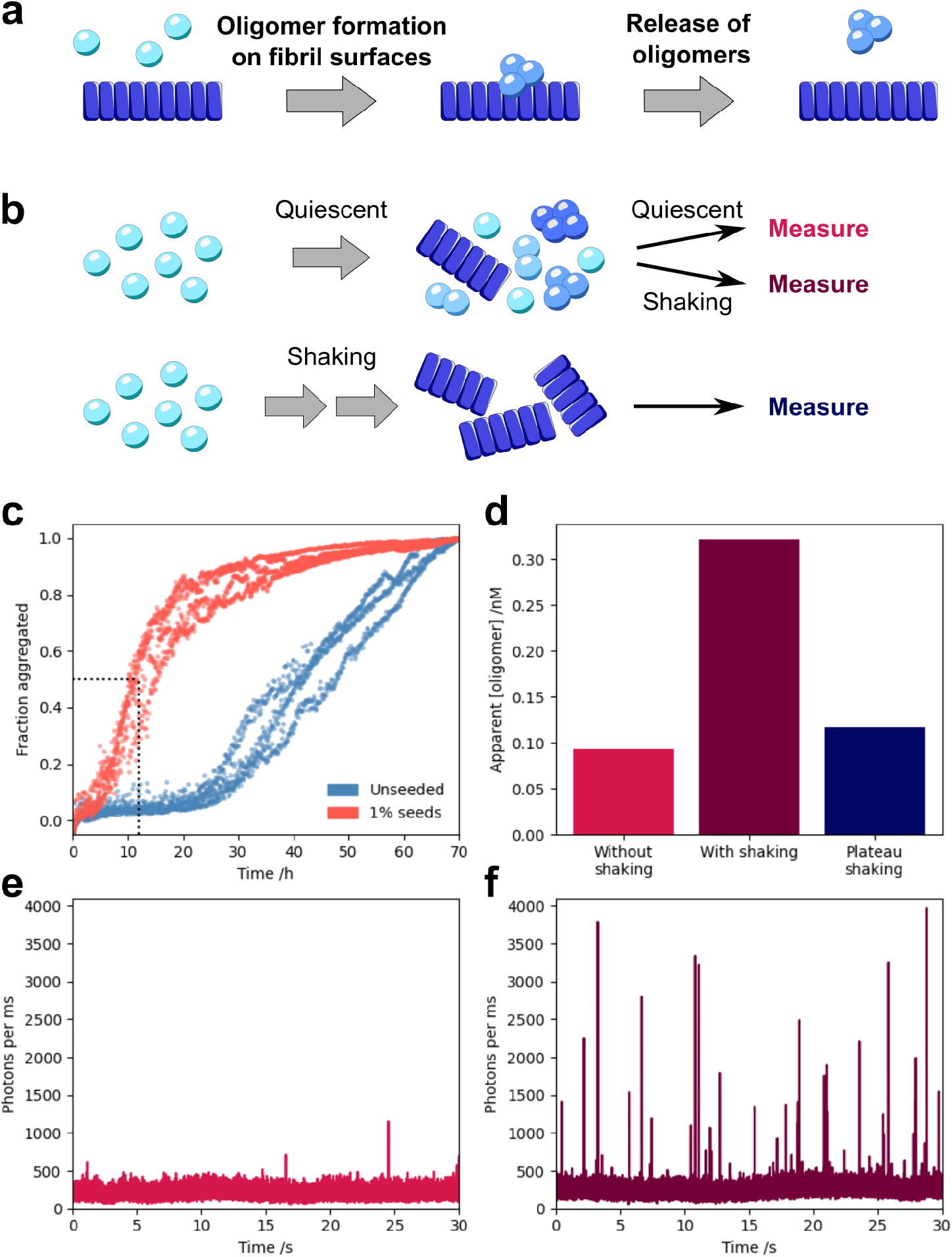
Secondary nucleation (**a**) proceeds under quiescent, physiological conditions. α-Synuclein was aggregated under both quiescent and shaking conditions (**b, c**), and the oligomer populations investigated. The relative oligomer mass concentrations of quiescent seeded (1%) aggregation reactions at the half-time before and after shaking for 10 minutes at 200 rpm are shown alongside the oligomer concentration during the plateau phase of the reaction under shaking (**d**). Example timetraces of photon count rates are shown for quiescent seeded (1%) aggregation reactions before (**e**) and after (**f**) shaking.

## Discussion

In this study, we have demonstrated that α-synuclein oligomers form by secondary nucleation under physiologically relevant conditions; namely neutral pH and a salt concentration mimicking the osmotic pressure in the cytosol. These oligomers are also the dominant mechanism of formation of new fibrils, allowing an exponential increase in aggregate mass concentration over time. Previous work has suggested that the contribution of secondary processes to α-synuclein aggregation is highly pH-dependent [34, 23, 21]. The presence of secondary processes at neutral pH was hinted at by data from Buell et al., and our detailed investigation we has now confirmed that the process is secondary nucleation and not fragmentation [21]. Moreover, several additional studies have reported seed-concentration dependent aggregation kinetics at neutral pH, suggesting that secondary nucleation is a general feature of α-synuclein aggregation [35, 36, 37, 34, 38].

Similarly, previous studies on α-synuclein oligomer dynamics have generally focused on the role of primary nucleation in oligomer formation [10, 17, 18]. In light of our findings herein, re-examination of these data shows that they are in fact consistent with a peak in concentration that we observe here, and thereby with a secondary nucleation formation mechanism. By investigating both fibril and oligomer dynamics in detail, we quantitatively elucidate the molecular mechanism of α-synuclein aggregation. This was made possible by our development of both aggregation conditions under cytosolic pH and ionic strength, and an oligomer detection method which is both blind to oligomer structure and minimally perturbs the reaction mixture. Through this study, we established that, while agitation is known to be able to induce fibril fragmentation, under these conditions it markedly increases the rate of secondary nucleation. This is analagous to crystallisation processes, for which mild agitation has been found to increase the rate of secondary nucleation by facilitating detachment, a finding which forms the basis of the use of mild agitation in industrial crystallisation processes [39, 40, 41, 42].

In conclusion, we identify secondary nucleation as the dominant process for the formation of both oligomers and fibrils in α-synuclein aggregation. α-Synuclein oligomers therefore provide both a potential source of toxicity and mechanism of aggregate spreading in PD [43, 7]. The detailed mechanistic framework we have elucidated can thus be used to better understand the role of α-synuclein in PD pathology and how to effectively develop therapeutic strategies.

## Acknowledgements

We thank Dr Manuela R Zimmermann and Minghao Zhang for helpful discussions on data analysis. We additionally thank Dr Heather Greer for her help with the acquisition of TEM images and the EPSRC Underpinning Multi-User Equipment Call (EP/P030467/1) for funding the TEM. We would like to acknowledge funding from the European Research Council under the European Union’s Horizon 2020 research and innovation program through the ERC grant DiProPhys (agreement ID 101001615), and the following sources: Herchel Smith Research Studentship (C.K.X.), Herchel Smith Fellowship (G.K.), Wolfson College Junior Research Fellowship (G.K.), Marie Sk-lodowska-Curie grant MicroSPARK (agreement no. 841466; G.K.), Swedish research Council (VR 2015-00143; S.L.).

## Contributions

C.K.X., G.K., W.E.A., M.V., S.L., and T.P.J.K. conceived the study. C.K.X. and M.C.C. developed the α-synuclein kinetics assay. C.K.X. and E.A. performed the kinetics experiments. C.K.X., E.A., and G.K. acquired µFFE data. C.K.X., G.M., and S.T. developed theory for data analysis. C.K.X., G.M., A.J.D., R.J., and W.E.A. contributed software. C.K.X., G.M., and A.J.D. analysed data. C.K.X. and G.M. wrote the manuscript with input from all authors.

## Notes

### Competing Interest Statement

The authors have declared no competing interest.

### Summary of Updates

Funding details in the acknowledgements section have been updated.

