## Supplementary information for "α-Synuclein oligomers form by secondary nucleation"

### Materials and methods

#### Purification of α-synuclein

α-Synuclein was purified as previously described [1]. α-Synuclein (WT or N122C variant) was overexpressed in Escherichia coli BL21 cells. The cells were centrifuged (20 min, 4000 rpm, 4 °C; JLA-8.1000 rotor, Beckman Avanti J25 centrifuge (Beckman Coulter)), and the pellet resuspended in buffer (10 mM tris, 1 mM EDTA, protease inhibitor) prior to lysis by sonication on ice. Debris was removed by centrifugation (20 min, 18,000 rpm, 4 °C; JLA-25.5 rotor), and the supernatant incubated (20 min, 95 °C). Heat-sensitive proteins were then removed by centrifugation (15 min, 18,000 rpm, 4 °C; JLA-25.5 rotor). Subsequent incubation with streptomycin sulphate (10 mg/mL, 15 min, 4 °C) precipitated out DNA. α-Synuclein was extracted from the supernatant (15 min, 18,000 rpm, 4 °C; JLA-25.5 rotor) by the gradual addition of ammonium sulphate (361 mg/mL). The α-synuclein-containing pellet was extracted by centrifugation (15 min, 18,000 rpm, 4 °C; JLA-25.5 rotor) and resuspended in buffer (25 mM tris, pH 7.4). Dialysis was used for complete buffer exchange, and the resultant mixture run at on a HiLoadTM 26/10 Q Sepharose high performance column (GE Healthcare), at room temperature. Under a gradient of 0-1.5 M NaCl over 600 mL, α-Synuclein was eluted at approximately 350 mM. Selected fractions were fractionated at room temperature on a Superdex 75 26/600 (GE Healthcare) and eluted in PBS (pH 7.4). For the N122C variant, 3 mM DTT was added to all buffers to prevent dimerisation. The The concentration of α-synuclein was determined by absorbance at 275 nm, with a molar extinction coefficient of 5600 $M^{-1}$ $cm^{-1}$. Aliquots were then flash-frozen in liquid nitrogen and stored at -80 °C.

#### Labelling of α-synuclein

The N122C variant of α-synuclein was fluorescently labelled as previously described [1]. DTT was removed from purified α-synuclein by buffer exchange into PBS using P10 desalting columns containing Sephadex G25 matrix (GE Healthcare). The DTT-free protein was incubated (overnight, 4 °C, rolling system) with a 1.5× molar excess of AlexaFluor-488 dye functionalised with a maleimide moiety. Excess unbound dye and α-synuclein dimers were removed by eluting the mixture over P10 desalting columns containing Sephadex G25 matrix (GE Healthcare). The resulting α-synuclein concentration was estimated by the dye absorbance at 495 nm, using a molar extinction coefficient of 72,000 $M^{-1}$ $cm^{-1}$. Aliquots were flash-frozen in liquid nitrogen and stored at -80 °C for up to 3 weeks prior to experiments.

#### Aggregation of α-synuclein

Aggregation of α-synuclein was carried out in non-binding 96-well plates (Corning) at 37 °C in a FLUOstar Omega microplate reader (BMG Labtech). Each well contained 100 µL of reaction mixture and a glass bead. The buffer used was Dulbecco's PBS (pH 7.4) with 0.01% (w/v) sodium azide. Interwell areas and empty wells were filled with PBS prior to sealing the plate with a foil cover. For experiments under shaking conditions, plates were shaken for 355 s at 200 rpm between each reading cycle; quiescent reactions were read at the same rate, but in the absence of all shaking. WT reactions were followed by the addition of 50 µM thioflavin T, whereas labelled N122C reactions (100% labelled protein) were monitored by AlexaFluor-488 fluorescence (Figure S7). Fibrils from unseeded reactions of 100 µM monomer were used directly as seeds for seeding reactions, with no sonication.

#### Analysis of bulk kinetic data

The signal obtained during aggregation kinetics (ThT fluorescence or AlexaFluor-488 fluorescence) was assumed to be proportional to the fibril mass present in the sample. The data were then fitted using the AmyloFit Platform and following the protocol in Meisl et al. to a model including primary nucleation, elongation and secondary nucleation with reaction order 0 [2]. This model was able to describe the data well across concentrations. To produce misfits, the same data were fitted with a model including only primary nucleation and elongation.

#### Measurement of fibril length distributions

At given timepoints in the plateau phase of the aggregation reaction, 1 µL of the reaction was withdrawn and the plate returned to the platereader. The reaction sample was mixed with 9 µL PBS (pH 7.4), and applied to a transmission electron microscopy (TEM) grid (continuous carbon film on 300 mesh Cu). Following adsorption, the sample was washed with milliQ water (2 x 10 µL). Samples were negatively stained with uranyl acetate (2% w/v, 10 µL, 2 min) and washed with milliQ water (2 x 10 µL). TEM grids were glow discharged using a Quorum Technologies GloQube instrument at a current of 25mA for 60s. TEM images were obtained using a Thermo Scientific (FEI Company) Talos F200X G2 microscope operated at 200 kV. TEM images were recorded using a Ceta 4k x 4k CMOS camera. The lengths of imaged fibrils were manually determined with ImageJ (example images in figure S9). The lengths of between 650 and 1550 individual fibrils were measured for each sample.

#### Analysis of fibril length distributions

The fibril length in the plateau phase of the reaction, during which the fibril mass concentration is constant, can be modelled by the following equations. By definition, in the plateau phase, the aggregate mass concentration is constant, i.e. $M(t) = M_\infty$. While nucleation processes become negligible when monomer is depleted, fragmentation still takes place, thus the number concentration of fibrils, $P(t)$, is given by

$$\frac{dP}{dt} = k_- M_\infty, \tag{1}$$

which can be solved to yield

$$P(t) = k_- M_\infty t + P_{plateau}, \tag{2}$$

where $k_-$ is the fragmentation rate constant, $t$ is the time since the plateau was first attained and $P_{plateau}$ is the number concentration of fibrils at $t = 0$.

The mean length, $L(t)$, at the plateau is thus given by:

$$L(t) = \frac{M(t)}{P(t)} = \frac{M_\infty}{k_- M_\infty t + P_{plateau}} = \frac{1}{k_- t + \frac{1}{L_{plateau}}}, \tag{3}$$

where $L_{plateau}$ is the average length when the plateau is first attained. Using the steady state expression for the average length during an aggregation reaction derived e.g. in Cohen et al. [3, 4, 5] as an estimate for $L_{plateau}$, the rate of fibril formation due to fragmentation is approximately given by $\kappa(\text{frag}) = L_{plateau} k_- = 0.01$ $h^{-1}$. This is thus an estimate of the rate of formation based purely on measurements of fibril lengths which can then be compared with kinetic measurements of the actual rate of fibril accumulation $\kappa$ to see if this is consistent with a purely fragmentation-driven mechanism.

#### Fabrication of microfluidic free-flow electrophoresis devices

Microfluidic free-flow electrophoresis (µFFE) devices were designed and fabricated as previously described [1]. Briefly, acetate masks were used to produce SU-8 moulds of devices by photolithography, the heights of which were measured with a profilometer (Dektak, Bruker). Polydimethylsiloxane (PDMS; 1:10 mixture of primer and base, Dow Corning) was applied to the mould and baked (65 °C, 1.5 h). µFFE devices were then excised and biopsy punches used to create holes for tubing and electrode connections, with diameters of 0.75 mm and 1.5 mm, respectively. Following sonication in isopropanol (5 min), devices were bonded to glass coverslips (#1.5) by activation with oxygen plasma. Immediately prior to use, prolonged oxygen plasma treatment was used to hydrophilise device surfaces.

#### µFFE device operation

The µFFE device design used contains liquid electrodes (3 M KCl solution containing 1 nM Atto-488 dye) to connect the electrophoresis chamber to the external electric circuit [6, 1]. These liquid electrodes were connected to the circuit via hollow metal electrodes made from bent syringe tips, which also constituted the outlets for the liquid electrodes. Samples were flowed into the device at controlled flow rates by the use of syringe pumps (Cetoni neMESYS, Korbussen, Germany), connected to polytetrafluoroethylene (PTFE) tubing (0.012" inner diameter × 0.030" outer diameter, Cole-Parmer, St. Neots, UK). The flow rates used were 1000, 200, 140, 10 µL $h^{-1}$ for the auxiliary buffer (15× diluted PBS in milliQ water), electrolyte, desalting milliQ water, and sample, respectively. The electric field was applied by a benchtop power supply (Elektro-Automatik EA-PS 9500-06, Viersen, Germany) connected to the metal electrode outlets.

#### Acquisition of µFFE data

µFFE data were acquired using laser confocal fluorescence microscopy; the microscope setup is described in detail in [7]. Briefly, a 488 nm wavelength laser beam (Cobolt 06-MLD 488 nm 200 mW diode laser, Cobolt, Stockholm, Sweden) was coupled into a single-mode optical fibre (P3-488PM-FC01, Thorlabs, Newton, NJ) and collimated (60FC-L-4-M100S-26, Schäfter und Kirchhoff, Hamburg, Germany) before being directed into the back aperture of an inverted microscope body (Applied Scientific Instrumentation Imaging, Eugene, OR). The laser beam was then reflected by a dichroic mirror (Di03-R488/561, Semrock, Rochester, NY) and focused to a concentric diffraction-limited spot in the microfluidic channel through a high-numerical-aperture water-immersion objective (CFI Plan Apochromat WT 60×, NA 1.2, Nikon, Tokyo, Japan). Photons arising thorugh fluorescence emission were detected using the same objective. Fluorescence was then passed through the dichroic mirror and imaged onto a 30 µm pinhole (Thorlabs), removing out of focus light. The signal was then filtered through a bandpass filter (FF01-520/35-25, Semrock), and focused onto a single-photon counting avalanche diode (APD, SPCM-14, PerkinElmer Optoelectronics, Waltham, MA). Photons were recorded using a time-correlated single photon counting (TCSPC) module (TimeHarp 260 PICO, PicoQuant, Berlin, Germany) with 25 ps time resolution. Single-photon counting recordings were obtained using custom-written Python code.

Aggregation samples (100 µL) were withdrawn from the plate at various times during the aggregation reaction and centrifuged (21,130 rcf, 10 min, 20 °C). The top 70 µL was carefully withdrawn without disturbing the pellet containing large aggregates. An aliquot of the supernatant was then diluted to approximately 5 µM total monomer mass concentration and injected into the device. An electric field of 300 V was applied and photon count timetraces obtained at 5-10 positions laterally distributed across the field direction, for a total of at least 1 min per position.

#### Analysis of µFFE data

Extraction of molecule concentrations and brightness from photon count data generally requires only one or a small number of molecules to simultaneously be located within the confocal spot and emitting photons. Such analysis approaches are therefore not suitable for our data, in which multiple protein species are always inside the confocal spot volume simultaneously. We therefore developed a theoretical framework which characterises the relationship between the detected distribution of photon count rates, and the brightness of the fluorophore. Our model comprises a confocal spot whose laser intensity is distributed as a 3-dimensional Gaussian function. Given that our data are acquired under laminar flow within a microfluidic device, and that the laser dimensions are approximately 10-fold smaller than the channel dimensions, our model entails that fluorophores are randomly distributed in the xy plane, and move in the z (flow) direction at the same speed, with no deviation in xy position (i.e. no diffusion on this timescale).

By integrating the total laser intensity that would be experienced at every possible xy position, we thus determined that the probability that a given fluorophore emits a total of $N^*$ photons while passing through the confocal volume is:

$$P_N(N = N^*) = \int_{I_{min}}^{I_{max}} \frac{e^{-alphaI}(\alpha I)^{N^*}}{IN^*!} dI \tag{4}$$

$$P_N(N = N^*) = \frac{\gamma(N^*, \alpha I_{min}) - \gamma(N^*, \alpha I_{max})}{N^*!} \tag{5}$$

where $I_{min}$ and $I_{max}$ are the minimum and maximum relative intensities of the laser regions through which the fluorophore is allowed to travel, and $\alpha$ is the expected number of total photons emitted when the fluorophore path goes through the centre of the laser. The probability that two non-interacting fluorophores which simultaneously flow through the confocal volume emit a total of $S^*$ photons is thus given by the convolution:

$$P_{S,2}(S = S^*) = (P_N * P_N)(N = N^*) \tag{6}$$

Similarly, the probability of $S^*$ total photons emitted from $n$ fluorophores can be determined by:

$$P_{S,n}(S = S^*) = (P_N * P_{S,n-1})(N = N^*) \tag{7}$$

With the assumption that the number of molecules simultaneously passing through the confocal volume is Poisson-distributed, the probability of detecting $D^*$ photons is:

$$P_D(D = D^*) = \sum_{n=0}^{\infty} \frac{\lambda^{M^*} e^{-\lambda}}{M^*!} P_{S,n}(S = D^*) \tag{8}$$

where $\lambda$ is the mean number of photons simultaneously located within the confocal volume. From plotting the expected distributions of photon counts arising from fluorophores at different concentrations and with different brightnesses, it is clear that these two parameters cannot compensate for each other; both parameters can therefore theoretically be determined from experimental data.

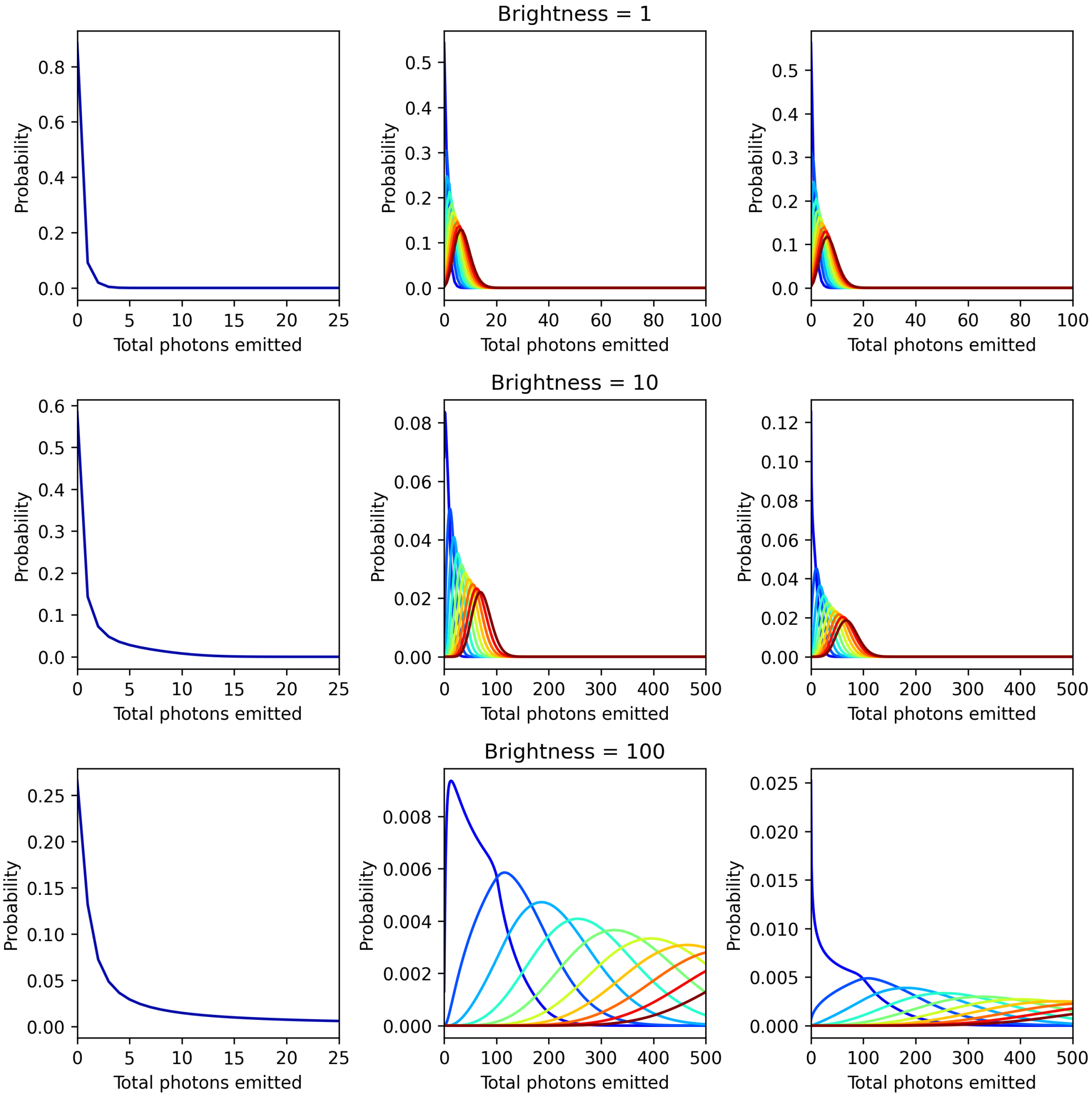


Figure S1: Expected photon count distributions of fluorophores passing through the confocal volume. Model distributions are shown for three different fluorophore brightnesses ($\alpha$ values of 1, 10, and 100). The expected distribution of emitted photons for a single fluorophore traversing the confocal volume (left) is shown alongside the expected distributions of multiple (5, 10, ... , 50 from blue to red) fluorophores being simultaneously located within the spot (middle). The right-hand plots show the expected distributions of emitted photons when multiple fluorophores pass through the confocal volume, with a mean of (5, 10, ... , 50 from blue to red) simultaneously inside the volume.

We fitted this model to data of AlexaFluor-488-labelled α-synuclein monomers (N122C variant) flowing through a straight microfluidic channel at different concentrations, with free parameters of concentration and brightness ($\alpha$). In all cases, the $\alpha$ value was fitted to be around 22 photons, and the relative concentrations differed by the expected factors. We note that the absolute concentration fitted is dependent on the dimensions of the confocal spot, which is not well defined; the values we took to be approximately the standard deviations of the laser intensity were determined by fluorescence correlation spectroscopy [7].

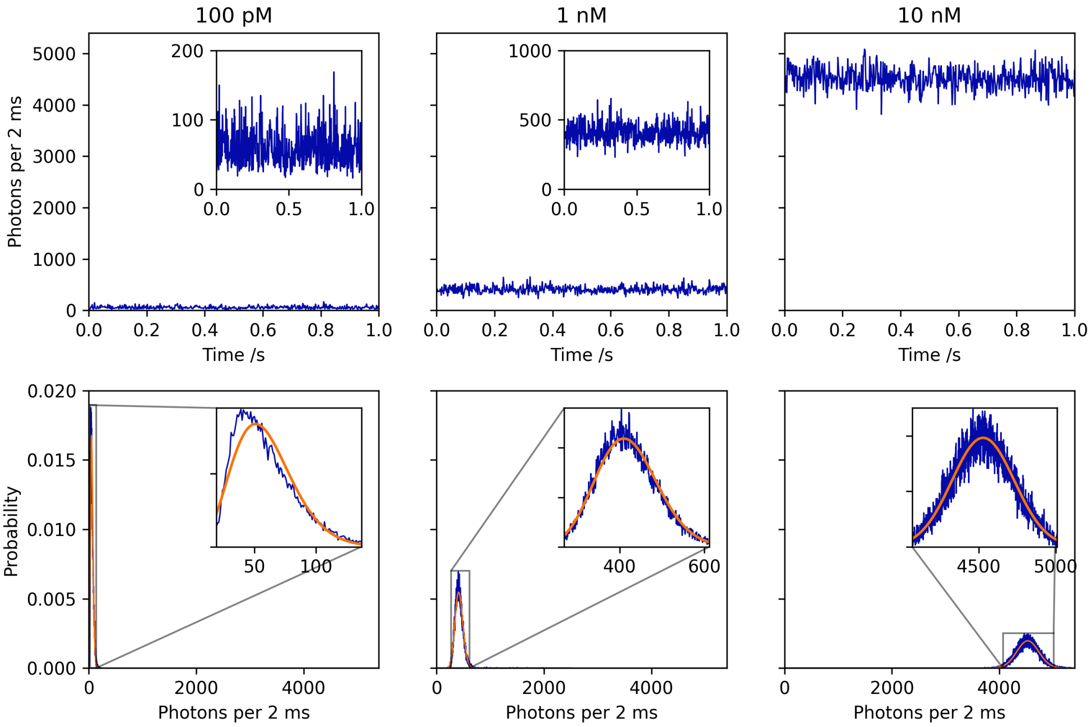


Figure S2: Determination of monomer brightness. Photon count data were acquired for monomeric α-synuclein at three different concentrations flowing through a straight microfluidic channel. Sections of the timetraces are shown in the top row, and their corresponding photon count distributions (blue) shown alongside the fitted distributions (orange) based on our model (equation 8) in the bottom row.

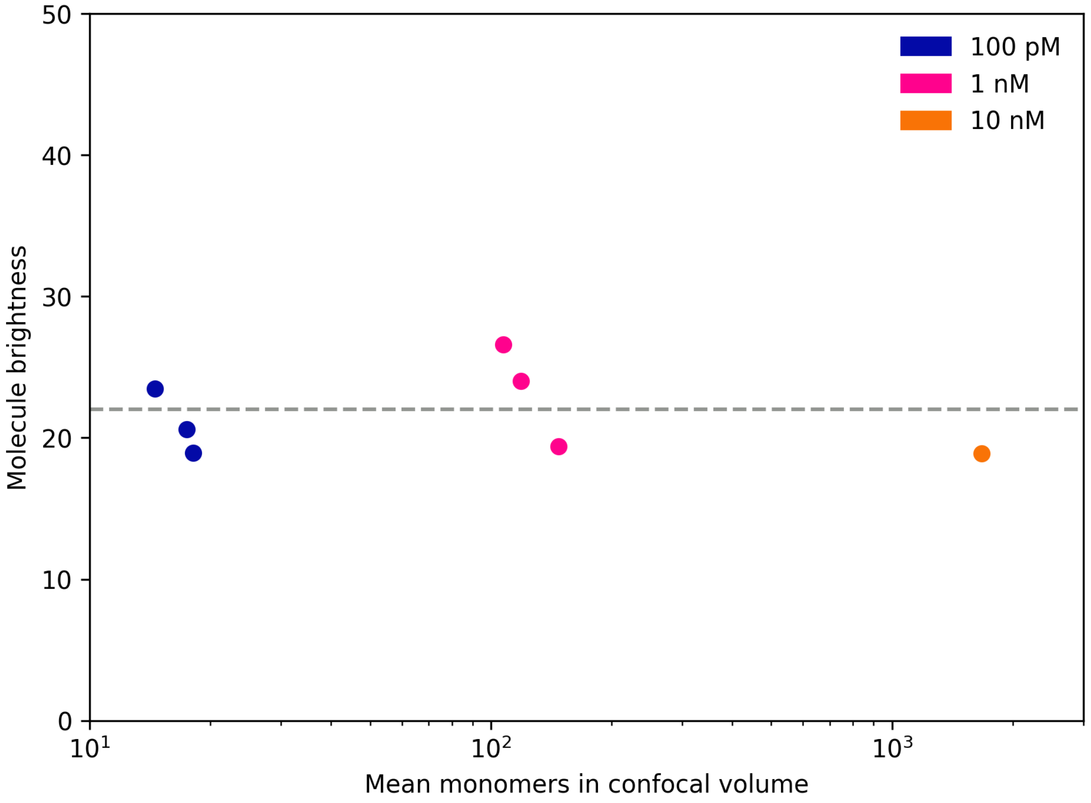


Figure S3: Fitted monomer parameters are consistent with experimental data. The fitted concentrations and brightnesses of labelled α-synuclein monomers are coloured according to the experimental concentrations. Both parameters are consistent across all data, validating our model. Based on these fits, the brightness value of monomers was taken to be 22 under our experimental conditions, as indicated by the grey dotted line.

Having determined the brightness of monomeric labelled α-synuclein, we next sought to determine the concentrations of oligomers from the photon count data. Oligomers contain multiple monomers and therefore multiple dyes, giving rise to a higher number of emitted photons per molecule; here, we estimate oligomer mass concentrations by assuming that dye molecules within oligomers contribute linearly to the total photons detected. In order to avoid potential artefacts introduced by flow or electric field instabilities, and to account for the background signal from monomers, the rolling median (50 ms window) was subtracted from the timetraces, before applying thresholds on the photon count rate to estimate the oligomer concentration.

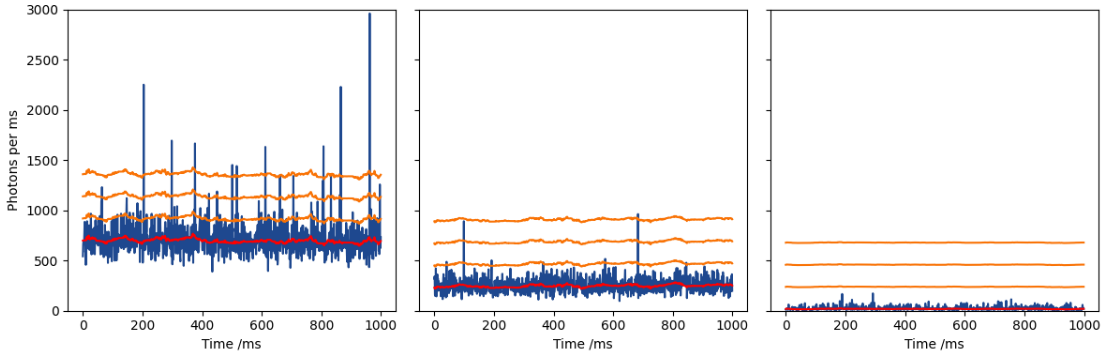


Figure S4: Example timetraces (blue) are shown alongside the rolling median (red) and oligomer thresholds (orange) of 220, 440, and 660 photons/ms, corresponding to 10-mers, 20-mers, and 30-mers, respectively. Example timetrace sections are shown with different oligomer contents.

However, this approach systematically underestimates the concentration of oligomers in the mixture. Using our model of photon count distributions detailed above, we calculated the fraction of oligomers which would be detected by using different photon count thresholds, for a range of oligomer sizes (Figure S5). Although the fraction detected is dependent on the size of the oligomers and threshold (440 photons in main text figure 4), the actual concentration of oligomers present in the aggregation mixture is likely to be at least tens to hundreds of fold higher than the apparent concentrations detected.

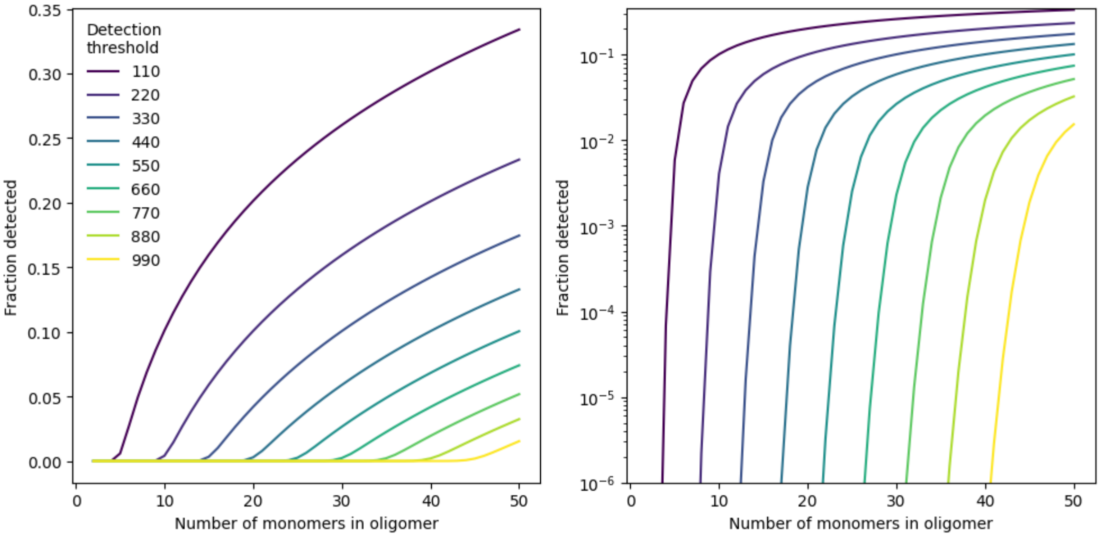


Figure S5: Fractions of oligomers that would be detected using different photon count thresholds.

In order to mitigate against variations in desalting efficiency and potential loss of protein by sticking to tubing, device, and/or syringe surfaces, the fraction of oligomer present in each timetrace was determined by comparing how many photon counts above the threshold compared to the total photon count. This fraction was then used in conjunction with the known supernatant concentration to obtain relative oligomer mass concentrations throughout the aggregation reaction.

#### Fitting of oligomer dynamics

The coarse-grained rate equations governing oligomers (concentration $S(t)$) formed by primary nucleation during an amyloid fibril formation reaction are:

$$\frac{dS}{dt} = k_{\mathrm{o}1} m(t)^{n_{\mathrm{o}1}} - k_{e1} S(t), \qquad (9)$$

where $m(t)$ is the concentration of monomeric protein. In a system dominated by secondary nucleation of oligomers, the rate equations are instead well-approximated by:

$$\frac{dS}{dt} = k_{\mathrm{o}2} m(t)^{n_{\mathrm{o}2}} M(t) - k_{e2} S(t) M(t), \qquad (10)$$

where $M(t)$ is the mass concentration of amyloid fibrils. For simplicity, we modelled $M(t)$ using the analytical expressions given in ref. [8] with rate parameters chosen as the values determined by fitting the ThT data on fibril formation in the main text. Eqs (9) and (10) were then fitted numerically to the experimental data on oligomer concentration using python. Eq. (10) provided the superior fit, supporting the conclusion that oligomers are formed predominantly by secondary processes in this assay.

### Supplementary figures

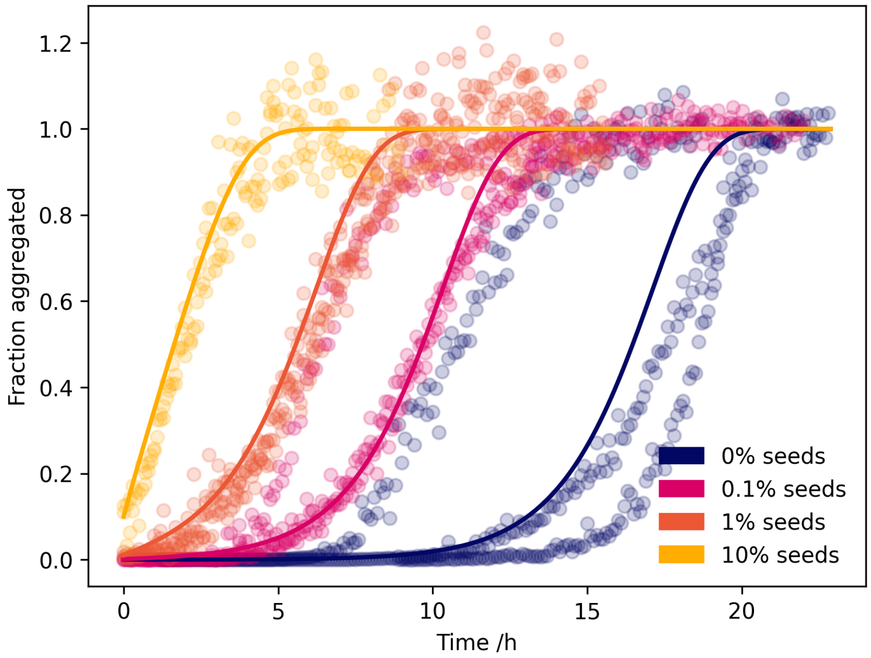


Figure S6: Labelling does not affect α-synuclein aggregation kinetics. Aggregation kinetics for unlabelled WT α-synuclein (100 µM total protein concentration) in the absence and presence of varying concentrations of seed fibrils, followed by thioflavin T fluorescence. Experimental data are shown as points, and fits as lines.

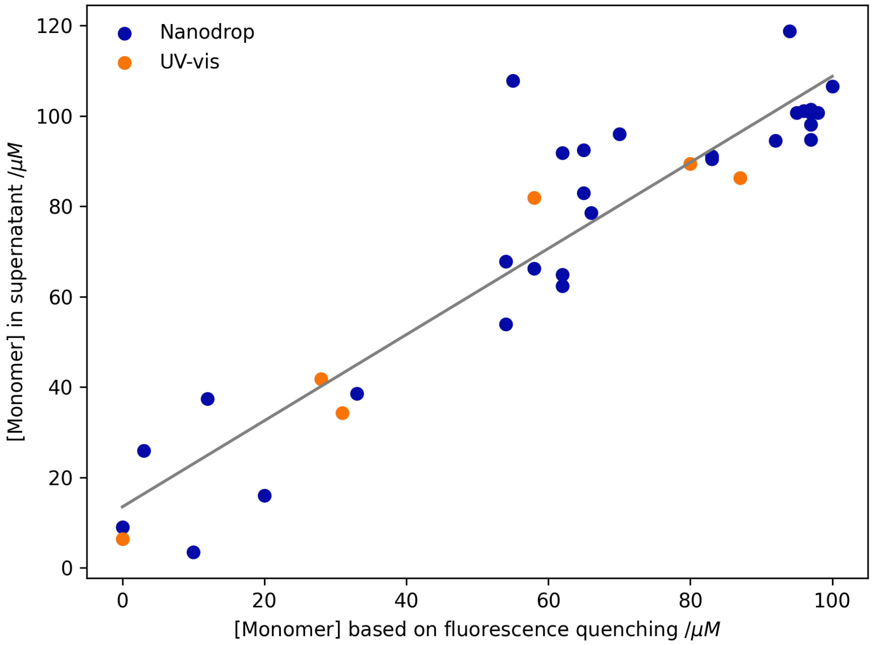


Figure S7: Aggregation of labelled α-synuclein can be followed by dye quenching. The concentration of the supernatant (21,130 rcf, 10 min) was determined by Nanodrop (blue) or UV-visible spectroscopy in a 1 cm-path length cuvette (orange), and compared to the estimated non-fibrillar concentration by fluorescence quenching measured in the platereader. The soluble α-synuclein concentrations from these complementary methods are in good agreeement, with the gradient of the line of best fit being 0.95, with an $R^2$ value of 0.93.

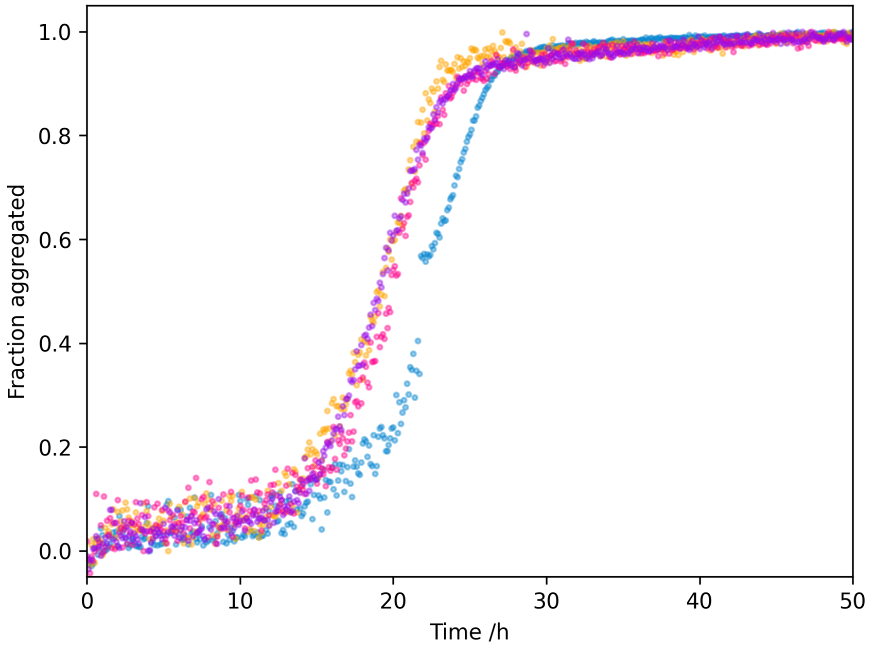


Figure S8: Example fibril formation kinetics of unseeded aggregation reactions of AlexaFluor-488-labelled N122C α-synuclein from different purification batches.

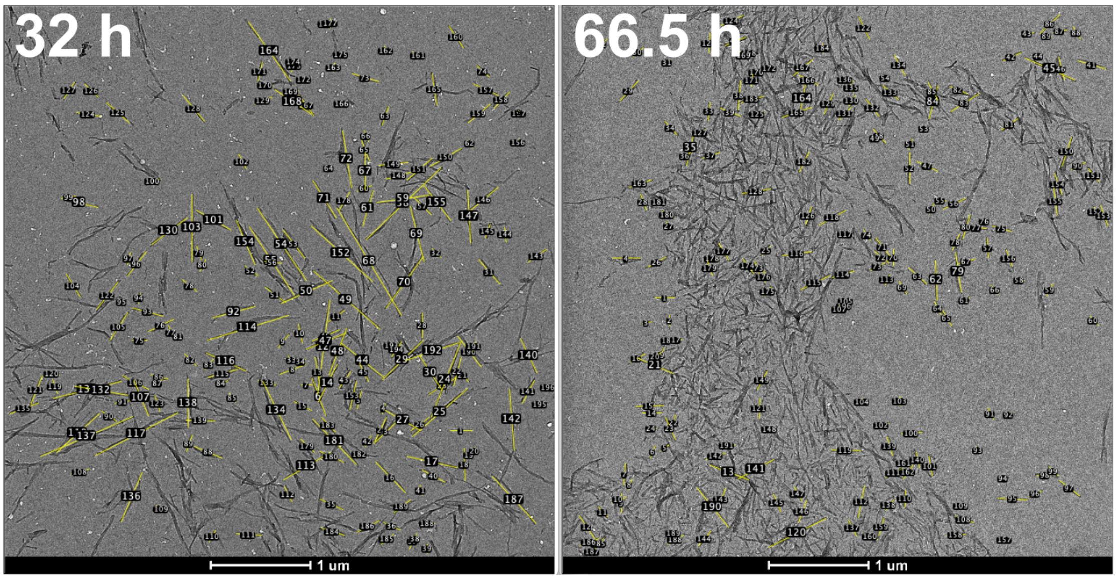


Figure S9: Example TEM images of α-synuclein fibrils. The lengths of fibrils where both ends were clearly visible were extracted using Fiji [9].

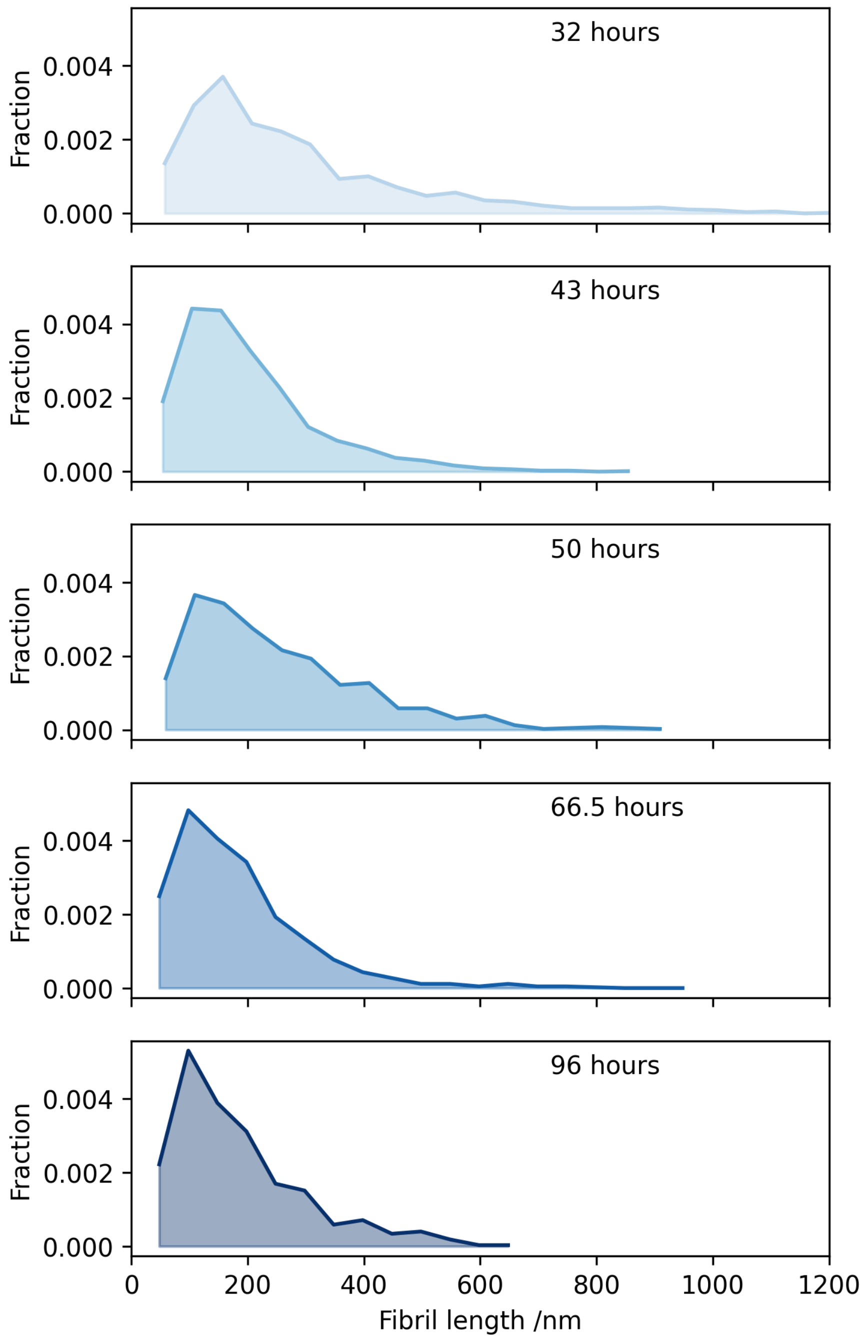


Figure S10: Length distributions of α-synuclein fibrils. Aliquots were extracted at various timepoints from an α-synuclein aggregation reaction during the plateau phase. Fibrils were imaged by TEM to determine their lengths.

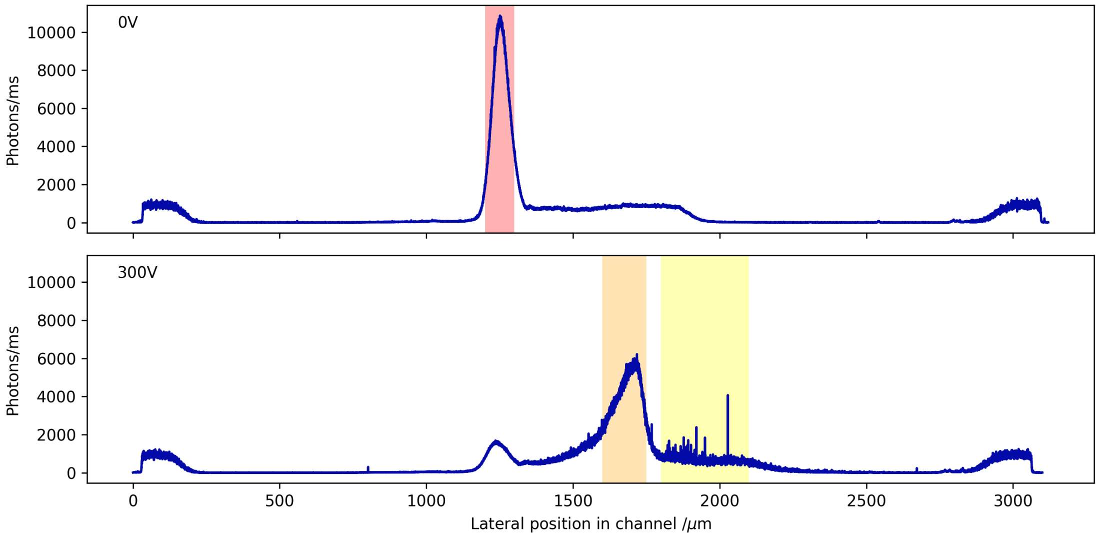


Figure S11: Oligomers have a higher mobility than monomers in the electric field. Examples of photon count rates across the channel lateral direction (parallel to the electric field). The edges of the channel are visible by the addition of Atto-488 dye to the electrolyte, evident here as the fluorescence peaks at the extremes of the channel positions. In the absence of the electric field, the whole aggregation mixture flows in a narrow stream (indicated by red). Upon the application of the 300V electric field, the sample is deflected laterally in the channel; the signal at the same position as in the 0V scan is due to protein stuck on the channel surface. Components can be separated by differences in electrophoretic mobility; the monomers (orange) are less mobile than the oligomers (yellow), due to the scaling of species charge and radius [1].
